# Cancer V-ATPase Expression Signatures: A Distinctive Balance of Subunit *C* Isoforms in Esophageal Carcinoma

**DOI:** 10.1101/489856

**Authors:** Juliana do Couto Vieira Carvalho dos Santos, Pedro Nicolau Neto, Evenilton Pessoa Costa, Frederico Firme Figueira, Tatiana de Almeida Simão, Anna Lvovna Okorokova Façanha, Luis Felipe Ribeiro Pinto, Arnoldo Rocha Façanha

**Affiliations:** Laboratório de Biologia Celular e Tecidual, Universidade Estadual do Norte Fluminense Darcy Ribeiro, Campos dos Goytacazes, RJ, Brazil; Programa de Carcinogênese Molecular, Instituto Nacional de Câncer - INCA, Rio de Janeiro, RJ, Brazil; Departamento de Bioquímica, Universidade Estadual do Rio de Janeiro, Rio de Janeiro, RJ, Brazil; Laboratório de Fisiologia e Bioquímica de Microrganismos, Universidade Estadual do Norte Fluminense Darcy Ribeiro, Campos dos Goytacazes, RJ, Brazil

**Keywords:** *ATP6V1C1*, *ATP6V1C2*, isoform *a*, cancer biomarker, esophageal cancer, V-ATPase

## Abstract

V-ATPases are hetero-oligomeric enzymes consisting of 14 subunits and playing key roles in ion homeostasis and signaling. Differential expressions of these proton pumps have been implicated in carcinogenesis and metastasis. To elucidate putative molecular signatures underlying these phenomena, we evaluated the V-ATPase genes expression in Esophageal Squamous Cell Carcinoma (ESCC) using gene expression microarray data and extended the analysis to other cancers the Oncomine database. Among all differentially expressed genes, those encoding the V-ATPase C isoforms exhibited striking expression patterns validated by qRT-PCR in paired ESCC samples and respective normal surrounding tissues. Structural modeling of C2a isoform uncovered motifs for oncogenic kinases in an additional peptide stretch, and an actin-biding domain downstream to this sequence. This study reveals multi-cancer molecular signatures in the V-ATPase structure and establishes that the expression ratios of its subunits/isoforms could form a conformational code that controls the pump regulation and interactions related to tumorigenic events.

## INTRODUCTION

Esophageal cancer (EC), the sixth most common cause of death from cancer worldwide (Bray, Ferlay et al. 2018, Ferlay, Colombet et al. 2018), encompass two histological subtypes, esophageal adenocarcinoma (EAC) and esophageal squamous cell carcinoma (ESCC), the last one responsible for about 90% of all diagnosed cases (Rustgi and El-Serag 2014). The very poor overall survival of ESCC patients is mainly due to its late stage detection and poor response to treatments (Bird-Lieberman and Fitzgerald 2009, Roshandel, Nourouzi et al. 2013). Therefore, it is essential to understand the molecular mechanisms involved in ESCC towards identifying new therapeutic targets and molecular biomarkers that can improve the detection and clinical managements.

The V-type H^+^-ATPase (V-ATPase) has attracted increasing attention in the field of molecular oncology (Stransky, Cotter et al. 2016) due to its differential overexpression in tumor cell membranes (Sennoune, Bakunts et al. 2004, Capecci and Forgac 2013) and the consequent intra- and extracellular pH dysregulation (Webb, Chimenti et al. 2011), both associated with key tumorigenic cellular processes, such as, migration and invasion (Sennoune, Bakunts et al. 2004, Capecci and Forgac 2013, Cotter, Capecci et al. 2015), regulation of signaling pathways frequently altered in cancer (Lim, Park et al. 2007, Cruciat, Ohkawara et al. 2010, Zoncu, Bar-Peled et al. 2011, Sennoune and Martinez-Zaguilan 2012, Faronato, Nguyen et al. 2015), immunomodulation (Katara, Jaiswal et al. 2014, Ibrahim, Kulshrestha et al. 2016) and therapy resistance (Martinez-Zaguilan, Raghunand et al. 1999, De Milito and Fais 2005, Liao, Hu et al. 2007, Wojtkowiak, Verduzco et al. 2011, Lu, Lu et al. 2013). These ATP-dependent proton pumps translocate H^+^ ions across endomembranes of all eukaryotic cells and also through plasma membranes of some specialized cells, acidifying intracellular compartments and the extracellular matrix, respectively (Nishi and Forgac 2002, Forgac 2007, Cipriano, Wang et al. 2008). The V-ATPase is a complex enzyme composed of 14 subunits organized into a cytoplasmic domain (V1) responsible for ATP hydrolysis and a proton translocation transmembrane domain (V0. The V1 domain consists of eight (A-H) subunits, while V0 sector, in human cells, is formed by the subunits *c, c”, a, d* and *e*, as well as two accessory subunits, *Ac45* and *M8-9* (Marshansky, Rubinstein et al. 2014, Cotter, Stransky et al. 2015).

The V-ATPase subunits *B*, *C*, *E*, *G*, *a*, *d*, and *e* have multiple isoforms encoded by different genes, which have been postulated to exhibit tissue specific expression patterns (Smith, Borthwick et al. 2002, Sun-Wada, Yoshimizu et al. 2003, Toei, Saum et al. 2010). However, up to now, no clear functional roles have been attributed to the distinct isoforms, except for the four *a* isoforms described as responsible for the enzyme targeting to different cellular membranes (Kawasaki-Nishi, Bowers et al. 2001), by which *a1* and *a2* are found in pumps of intracellular membranes, while pumps assembled with *a3* and *a4* are usually directed to plasma membrane (Morel, Dedieu et al. 2003, Toyomura, Murata et al. 2003, Wagner, Finberg et al. 2004, Toei, Saum et al. 2010, Capecci and Forgac 2013). In addition, it has also been demonstrated that yeast V-ATPase complexes containing two different *a* isoforms show distinct cellular localization, coupling efficiency and regulation by association/dissociation of the V1 and V0 domains (Kawasaki-Nishi, Nishi et al. 2001, Samarao, Teodoro et al. 2009]). V-ATPase plays essential roles in a variety of cellular processes, and the existence of multiple subunit isoforms might reflect multifunctional conformations, including specific compositions for specific cell types and pathophysiological conditions. Here, we investigated the differential expression patterns of the V-ATPase subunits/isoforms in human ESCC and in other types of tumors, exploring the potential of these proton pumps as putative cancer biomarkers for early diagnosis and more effective prognosis.

## RESULTS

### Expression profile of V-ATPase subunits in ESCC

The expression profiles of 25 genes encoding V-ATPase subunits were obtained from an ESCC transcriptome developed by the Brazilian National Institute of Cancer – INCA (Nicolau-Neto, Da Costa et al. 2018). An unsupervised hierarchical clustering analysis showed that ESCC exhibits striking changes in expression of most isoforms of the V-ATPase subunits in comparison with normal esophageal tissues excised surrounding the tumors, in a profile which clearly segregates these two groups and suggests a pattern of tumor-specific V-ATPase expression (**Figure. 1a**; the fold change and *p*-value of each gene is shown in **Table S1**). The genes were considered differentially expressed when their *p*-value was less than 0.005. It is of note that the *A* and *B* subunits, which constitute the evolutionary conserved catalytic domain of the V-ATPase, presented no significant expression changes (*ATP6V1A*: 1-fold change and p = 0.78; *ATP6V1B1*: 1-fold change and p = 0.79; *ATP6V1B2*: −1.08-fold change and p = 0.45) (**Table 1**, **Figure S1**).

**Figure 1.**
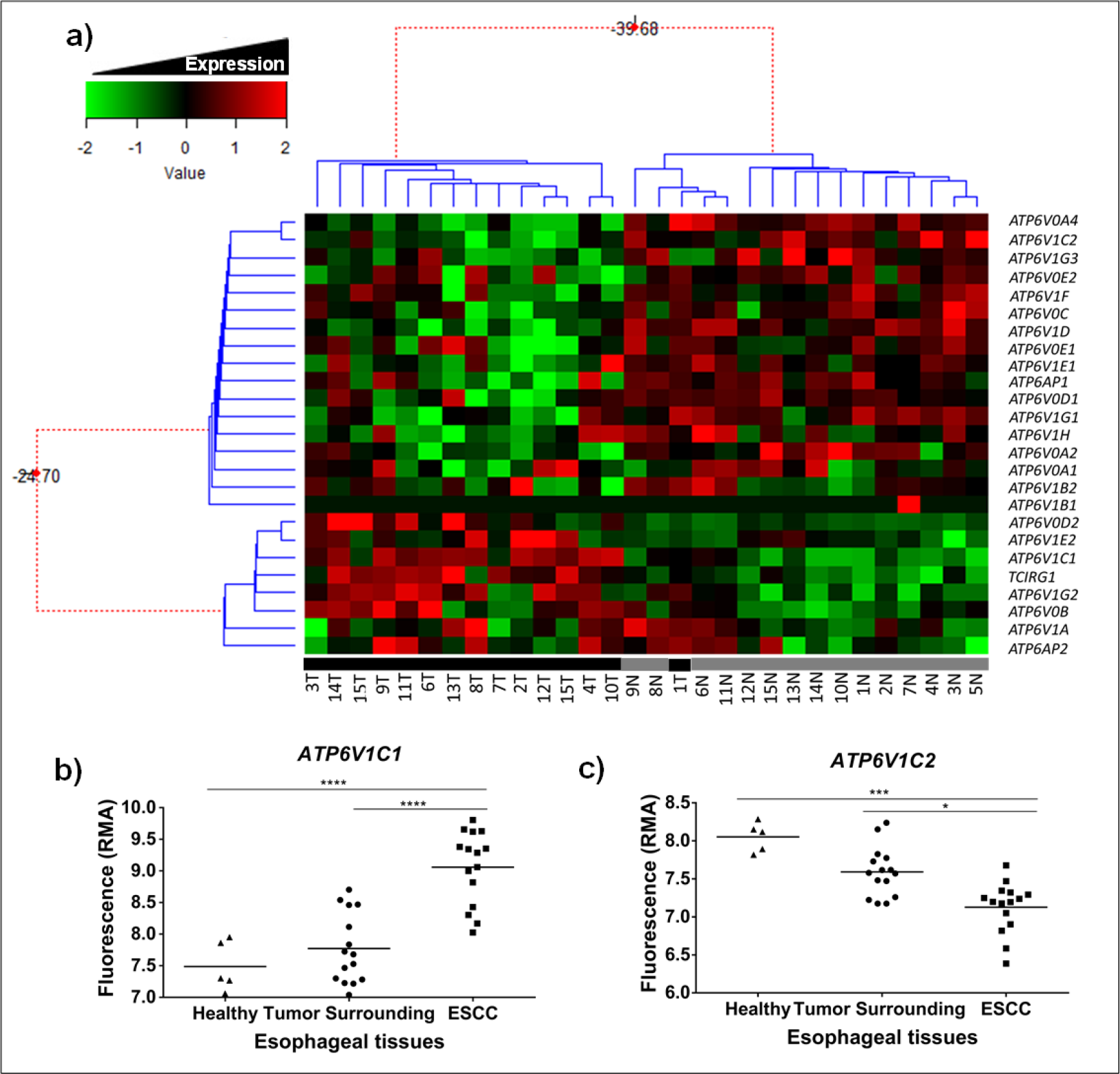
V-ATPase expression profile in ESCC by microarray database. (a) Unsupervised hierarchical clustering analysis of 15 esophageal samples (tumors and respective surrounding tissues) showing the expression of the 25 genes of V-ATPase. Green indicates decreased gene expression and red increased genes expression. N and black bar = normal surrounding tissue; T and grey bar = Tumor. (b) *ATP6V1C1* and *ATP6V1C2* expression levels in tumor, normal-appearing surrounding tissue and healthy esophageal mucosa from individuals without cancer. *p < 0.05; ***p < 0.001; ****p < 0.0001.

The subunit *c”* (*ATP6V0B*), which compose the proteolipid ring of the V-ATPase, exhibited higher expression in ESCC samples than in normal tissues. Even more interesting were the changes found among the isoforms expression of the same subunit, as observed for *d* and *G* subunits, where, *d2* (*ATP6V0D2)* and *G2* (*ATP6V1G2)* isoforms were more expressed in tumor, while *d1*, *G1* and *G3* isoforms (*ATP6V0D1, ATP6V1G1* and *ATP6V1G3*, respectively) were less expressed in tumor compared with normal surrounding tissue (**Figure S2**).

However, among all differentially expressed V-ATPase genes, only genes for the subunits *C* and *a* presented the highest differences among their respective isoforms. In ESCC compared with normal surrounding tissue, *ATP6V1C1* and *TCIRG1* mRNAs were upregulated (2.79 and 1.53-fold change, respectively), while those of *ATP6V1C2* and *ATP6V0A4* were downregulated (−1.28 and −2.76-fold change, respectively). In addition, significant differences in expression in these four genes were also observed between ESCC and healthy esophageal tissues from individuals without cancer (**Figure 1b** and **c**; **Figure S3**). The *TCIRG1* and *ATP6V0A4* genes encode two of the four distinct *a* isoforms, *a3* and *a4* (V0 sector), whereas in the V_1_ domain, *ATP6V1C1* and *ATP6V1C2* encode the different *C* subunit isoforms; *ATP6V1C1* encodes the *C1* isoform (NCBI gene ID: 528) and *ATP6V1C2* encodes two alternative transcriptional splice variants, namely *C2a* and *C2b* isoforms (NCBI gene ID: 245973).

To investigate the expression pattern of V-ATPase genes in others cancer types, we systematically analyzed mRNA transcriptome between normal tissues and tumor tissues of 18 sites comprising different histological types using the Oncomine database. The threshold was designated according to the following values: *p*-value 1E-4, fold-change 2, and top gene ranks 10%. Was possible to verify a common regulation of the V-ATPase isoforms expression in different tumor types. Remarkably, *ATP6V1C1* and *TCIRG1* expression was constantly higher in most cancer types compared to normal tissues, while *ATP6V1C2* and *ATP6V0A4* were downregulated, in the same way as founding the ESCC microarray experiments (**Figure S4**). These results imply a coordinate expression and a balance between specific isoforms of the subunits *C* and *a* in tumors.

### *ATP6V1C1* and *ATP6V1C2* transcripts are differentially expressed in ESCC

In order to validate the most expressive results of the microarray analysis, we evaluated *ATP6V1C1* and *ATP6V1C2* mRNA levels by RT-PCR in 38 ESCC paired samples (tumors and respective normal surrounding tissues). The *ATP6V1C1* expression in ESCC relative to respective normal tissue ranged from 0.43 to 7.37-fold change, with the median value of 2.33-fold change. Assuming a fold change cut-off of ≥ 2, about half (54%) of the ESCC samples presented *ATP6V1C1* mRNA levels increased when compared to their respective normal surrounding tissues (**Figure 2a**), while the expression of the *ATP6V1C2* gene was downregulated in most of the tumors analyzed, considering a cut-off ≤ −2 (67.6% for v1; 72.2% for v1,2). For *ATP6V1C2* fold change values ranged from −0.46 to −48.9 with the median value of −3.96 for *v1* and of −0.51 to −52.9 with the median value of −6.53 for *v1,2* (**Figure 2b** and **c**).

**Figure 2.**
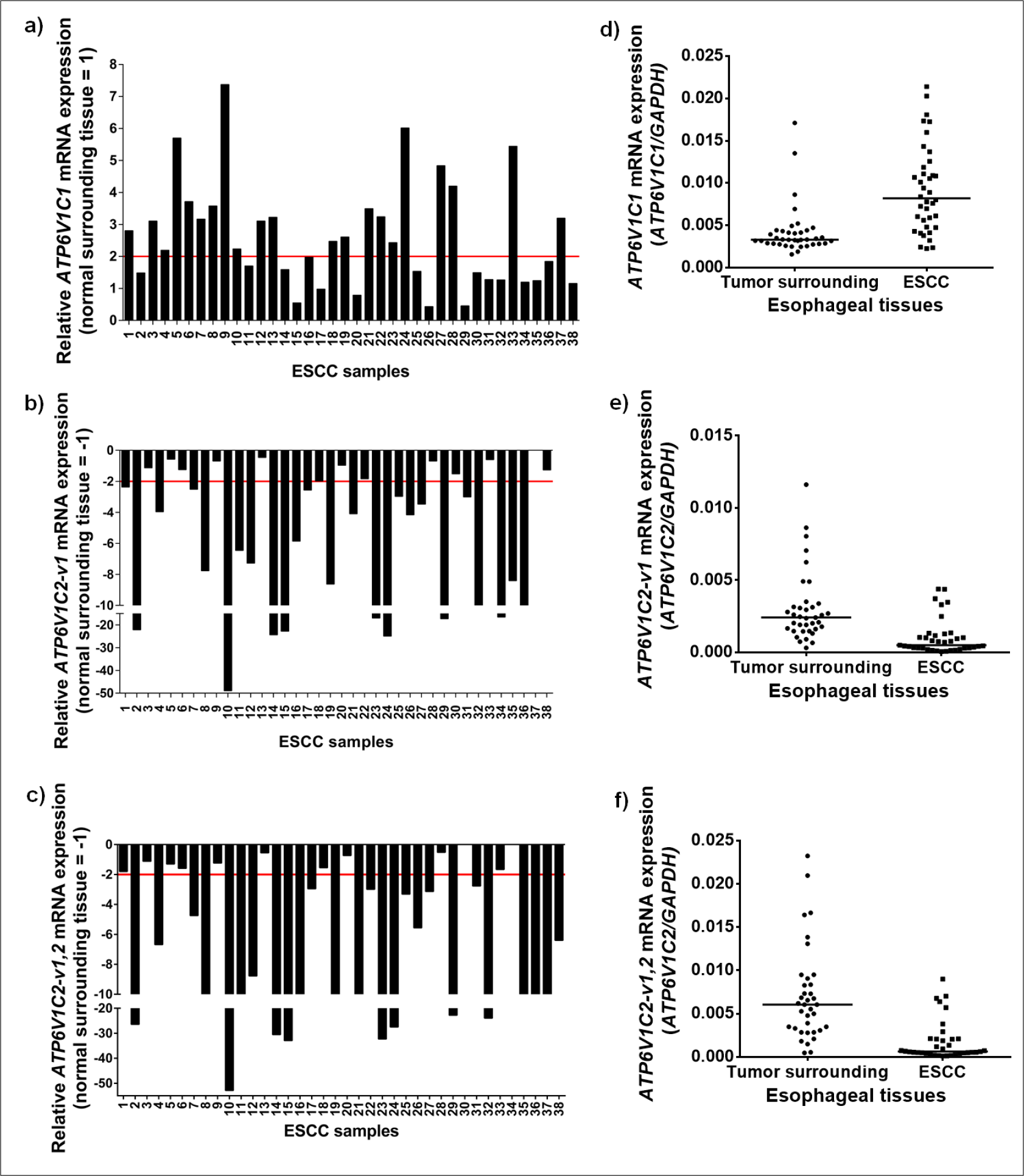
The expression pattern of V-ATPase *C* isoforms in ESCC. *Left images:* RT-PCR analysis of (a) *ATP6V1C1*, (b) *ATP6V1C2*-v1 and (c) *ATP6V1C2*-v1,2 mRNA levels in paired ESCC samples. ESCC samples presenting over 2-fold increase (red dashed line) in relative *ATP6V1C1* expression were considered upregulated and ESCC samples presenting decrease of −2-fold (red dashed line) in relative *ATP6V1C2* expression were considered downregulated. Values are expressed as relative to those obtained in tumors respective normal surrounding tissue (= 1 or −1). *Right images:* qRT-PCR evaluation of (d) *ATP6V1C1*, (e) *ATP6V1C2*-v1 and (f) *ATP6V1C2*-v1, 2 mRNA levels distribution in the groups of normal surrounding and their paired ESCC tissues. Values are shown in relative units. mRNA levels were normalized by those of *GAPDH*, used as the housekeeping gene. p < 0.0001.

Next, a comparative analysis of mRNA levels of subunit *C* isoforms in tumors and normal surrounding tissues was performed. Expression levels of *ATP6V1C1* in ESCC samples group was approximately 2.5-fold higher than those detected in the normal surrounding tissues group (median values of 0.008196 and 0.003301, respectively) (**Figure 2d**). For *ATP6V1C2-v1* and *ATP6V1C2-v1,2* the mRNA levels were significantly lower in the ESCC group than those in the normal surrounding tissues group, with the median values of −4.3 and −9.6-fold, respectively (*v1*: median of 0.000489 in ESCC and 0.00241074 normal surrounding; *v1,2*: median of in 0.000629 ESCC and 0.006044 normal surrounding) (**Figure. 2e** and **2f**). These findings are in agreement with the data obtained from microarray analysis (**Figure 1**), confirming that the V-ATPase *C* isoforms present a remarkable differential expression in ESCC, when compared to normal surrounding tissue.

We further examined the expression ratio of *ATP6V1C1* and *ATP6V1C2* isoforms (*C1*/*C2-v1 and C1*/*C2-v1,2* ratios). The median value of *C1*/*C2-v1* ratio was of 1.4-fold in normal surrounding tissues and 13.4-fold in the ESCC samples, while the median value of *C1*/*C2-v1,2* ratio was of 0.5-fold and 10.1-fold in the normal surrounding and ESCC tissues, respectively (**Figure S5**). These results suggest that the expression levels of the different *C* isoforms are carefully guarded to achieve a proportional balance in normal tissues, which dysregulation towards the *C1* prevalence over the *C2* is an intrinsic characteristic of ESCC.

### *C* isoforms as potential diagnostic biomarker for ESCC

In order to evaluate whether *C* isoforms mRNA expression could be used to discriminate between tumor and non-tumor esophageal tissues, we performed the Receiver Operating Characteristic (ROC) curve analysis using the gene expression values from ESCC and normal surrounding tissues. The differential expression of *ATP6V1C1* and *ATP6V1C2* was able to accurately discriminate ESCC samples from normal surrounding tissues (p<0.0001), with sensitivity and specificity of 79.49% and 84.62% for *ATP6V1C1;* 83.78% and 83.78% for *ATP6V1C2-v1* and 80.56% and 86.11% for *ATP6V1C2-v1,2*, respectively. The ROC curve analysis for the *C1*/*C2* expression ratios generated an area under curve greater than 0.94, reflecting the high accuracy of the test to discriminate tumors and normal surrounding tissues. The *C1/C2-v1* and *C1/C2-v1,2* ratios showed respectively a sensitivity of 86.49% and 88.89% and a specificity of 91.89% and 91.67% (**Figure 3**). These results reveal that mRNA expression ratio of the subunit *C* isoforms can be useful as a diagnostic biomarker for ESCC.

**Figure 3.**
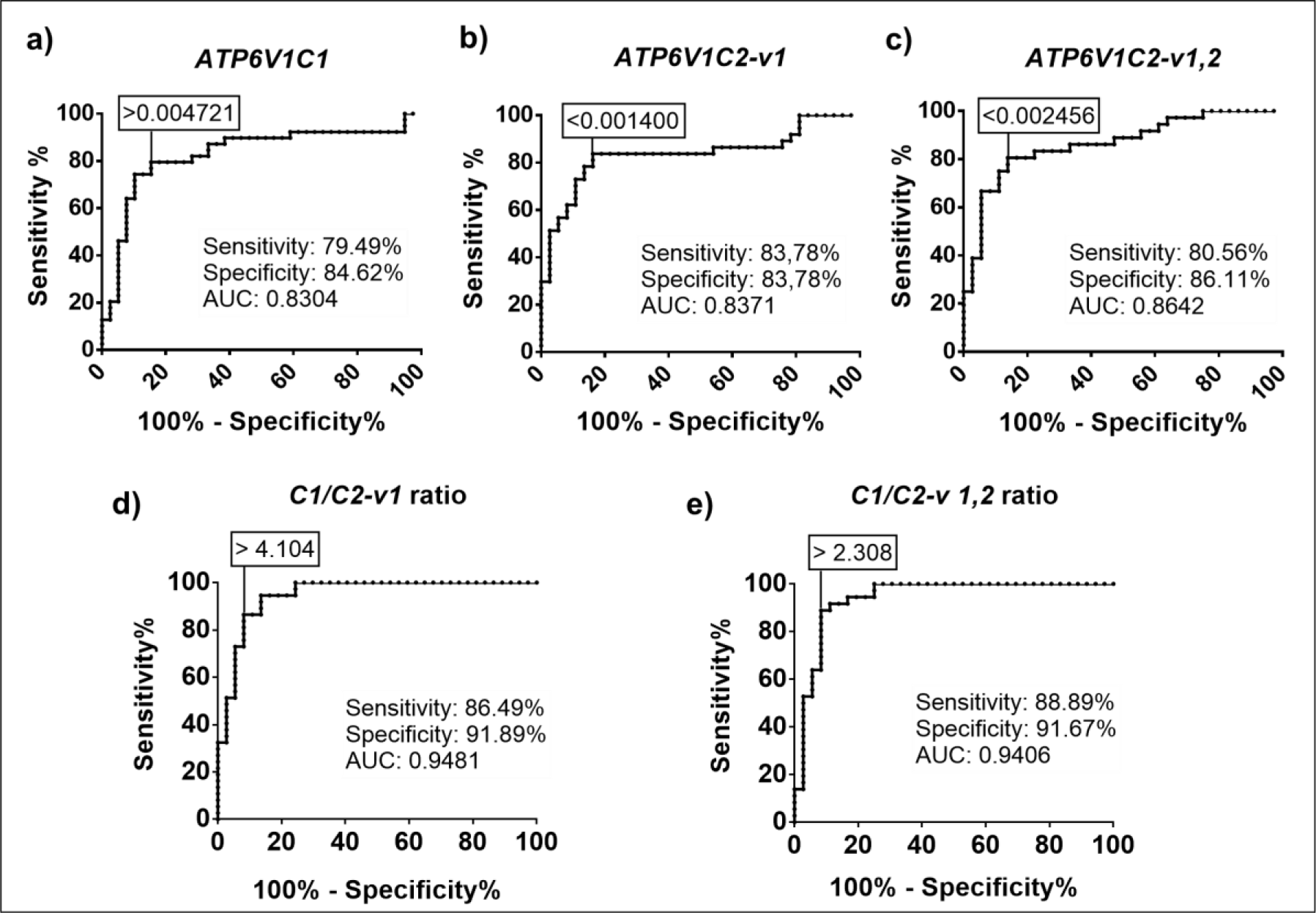
ROC curves of C isoforms mRNA expression for the discrimination of ESCC tissue from normal surrounding mucosa samples. ROC curves are relative to mRNA expression of: (a) *ATP6V1C1*, (b) *ATP6V1C2*-v1, (c) *ATP6V1C2*-v1,2, (d) *C1*/*C2*-v1 ratio and (e) *C1*/*C2*-v1,2 ratio. The area under curve (AUC) indicates the accuracy of the test in discriminating tumors from normal surrounding tissues. Numbers in a box correspond to a cut-off used for the determination of the sensitivity and specificity. ROC, receiver operating characteristic. p < 0.0001.

### Expression profiles of V-ATPase genes in EC histological subtypes

The expression profile of two genes of the V-ATPase subunit C in ESCC prompted us to investigate if such dysregulation is involved in esophageal carcinogenesis in general, independently of histological subtype. To this end, we used TCGA database to analyze the expression levels of V-ATPase genes in ESCC and EAC. The expression profile of all V-ATPase genes presented a very similar pattern in both histological subtypes (**Figure 4a**). Regarding to C isoforms, both EAC and ESCC tumors exhibited higher expression levels of *ATP6V1C1* than *ATP6V1C2*. The *ATP6V1C1* expression was increased 6.6-fold when compared to *ATP6V1C2* expression in EAC tissues. In ESCC samples, *ATP6V1C1* was 11.6-fold more expressed than *ATP6V1C2*, corroborating with the data obtained by qRT-PCR, where *ATP6V1C1* was 12.7-fold more expressed in tumors than *ATP6V1C2* (**Figure 4b**). Finally, the *ATP6V1C1* expression levels were similar between the two-histological subtypes (expression median of 1578 for ESCC and 1479 for EAC), however, the average expression of *ATP6V1C2* was lower in ESCC than in EAC (expression median of 136 for ESCC and 225 for EAC) (**Figure S6**).

**Figure 4.**
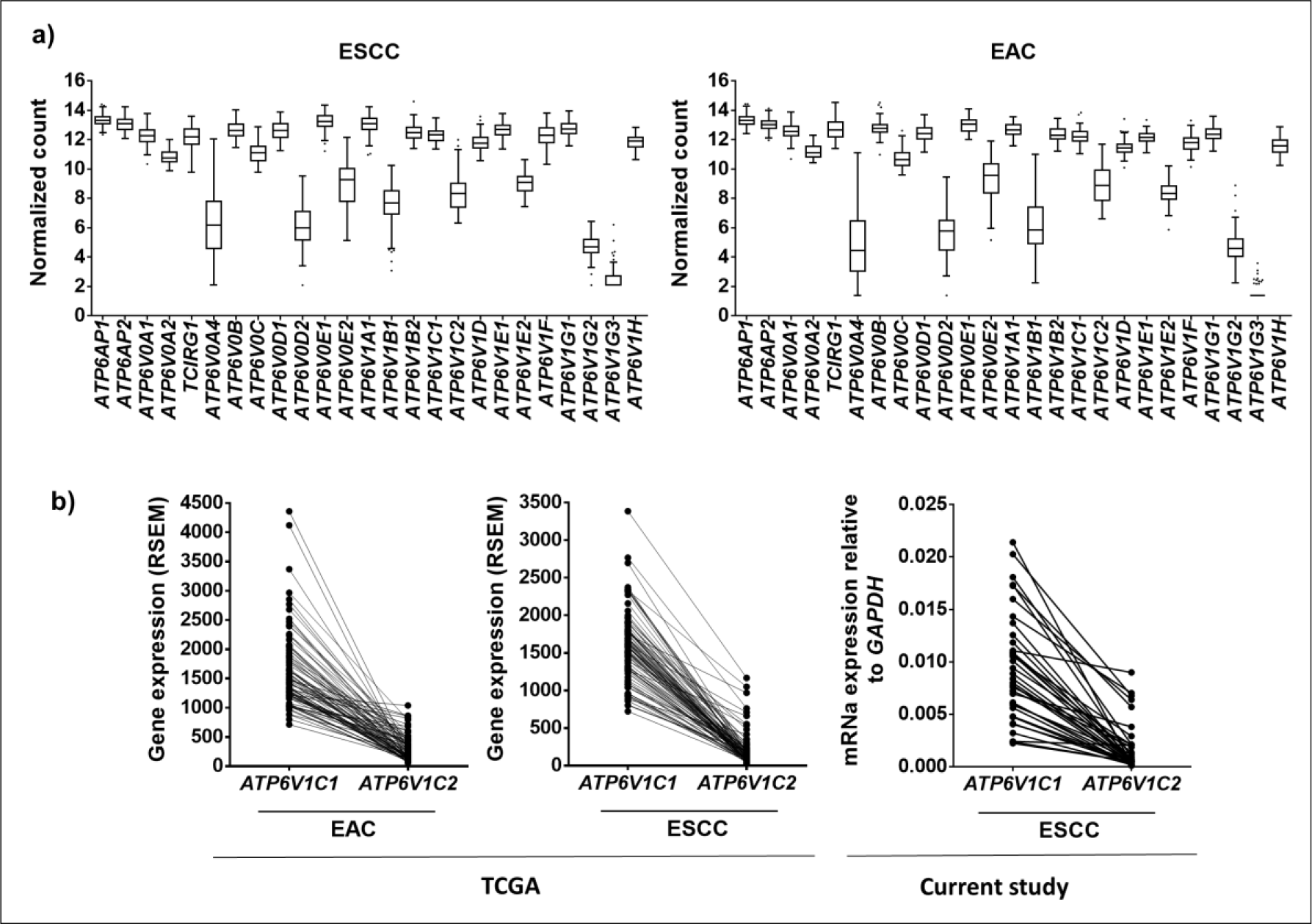
V-ATPase mRNA expression in esophageal cancer histological subtypes. (a) Expression profile of all V-ATPase genes in esophageal adenocarcinoma (EAC) and ESCC. (b) Respective *ATP6V1C1* and *ATP6V1C2* mRNA levels in EAC and ESCC assessed by RNA-sequencing data from TCGA and by qRT-PCR in the current study.

### Structural modeling of the subunit C isoforms

In addition, we performed the analysis of conserved domains among the human V-ATPase subunits *C1* (NP_001686.1) (hC1), *C2a* (NP_001034451.1) (hC2a) and *C2b* (NP_653184.2) (hC2b) using the program BLASTP 2.8.0 in the mammalian taxon of the database *NCBI Protein Reference Sequences*. The three amino acid (aa) sequences were found to be highly conserved, and hC1 (382 aa), hC2a (427 aa) and hC2b (381 aa) share at least ~55,3% identity and ~74,2% similarity (**Table 2**). When compared to isoform hC2b, the isoform hC2a has an additional exon that encodes for 46 aa at the position 276-321 aa. The BlastP analysis suggested that the isoform hC2a is present in all mammals, but absent in other taxa (yeast, nematode, mollusk, arthropod, birds, reptiles, amphibia and higher plants). Next, were performed analysis for protein-protein binding and for phosphorylation sites, exclusively on hC2a additional exon. These analyses revealed interesting motifs for serine/threonine kinases and phosphatases, such as ERK1 and 2, Casein Kinase II, PKA, PKC; and three potential binding motifs for proteins which recognize phosphorylated serine-based motifs with binding domains to WW, MDC1 BRCT and Plk1 PBD (**Tables 3** and **4**). No tyrosine kinase/phosphatase binding sites were identified in this specific region. Predictions of hC1, hC2a and hC2b three-dimensional structures were performed by homology modeling and *ab initio* approaches (**Figure 5**). The RMSD assessment also indicated that hC1 and hC2b are structurally very similar, while hC2a has an additional structural motif (**Table 2**). Analysis of global and local stereochemical quality using validation programs revealed that the most stable structure predicted for the additional exon of hC2a would be an alpha-helix conformation flanked by unstructured regions (**Figure 5c**). However, since the subunit *C* binds to, and serves as a substrate for PKA and PKC (Voss, Vitavska et al. 2007), these proteins were also used to refine the validation of the hC2a structural model. Following a series of protein-protein dockings between proposed hC2a models with PKA and PKC structures (**Figure S7**), the conformational model that best established PKA/PKC-hC2a interaction complexes was the hC2a predicted model for which the domain corresponding to the additional exon is in a beta-sheet flanked by unstructured regions (**Figure 5d**, **Figure S7**).

**Figure 5.**
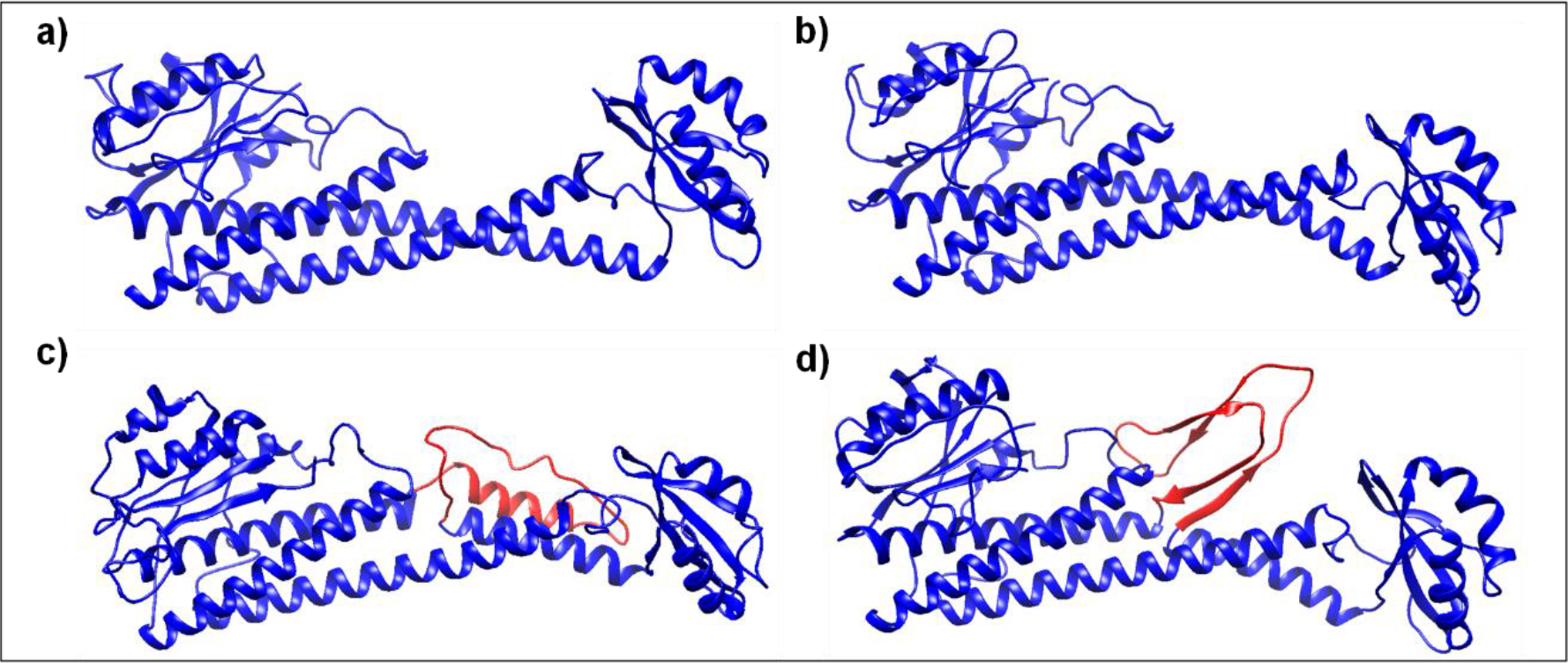
Prediction of three-dimensional structure of *C* subunit isoforms. (a) hC1 isoform, (b) hC2b isoform, (c) most promising model of hC2a isoform and (d) model of hC2a isoform validated by protein docking. C1 and C2b were modeled by homology using as template the resolved structure from yeast V-ATPase (3J9V.pdb) and C2a was obtained by *ab initio* approach. The additional peptide stretch present in the hC2a isoform is shown in red.

## DISCUSSION

In recent years, there has been an increasing body of evidence establishing that the V-ATPase proton pump is implicated in several cancer hallmarks (Hanahan and Weinberg 2011), such as, resistance to cell death, increased cell proliferation, invasion and metastasis, induction of angiogenesis, immune system modulation and reprogramming of energy metabolism (Webb, Chimenti et al. 2011, Stransky, Cotter et al. 2016, Pamarthy, Kulshrestha et al. 2018). However, despite the large amount of data accumulated on V-ATPase differential expression in various types of tumors, no clear molecular signature could be related to the carcinogenesis process in general. In previous reports, the overexpression of different subunits and isoforms have been highlighted in different cancers (Whitton, Okamoto et al. 2018), including a3 isoform in ovarian cancer (Kulshrestha, Katara et al. 2015), C1 isoform in oral cancer (Otero-Rey, Somoza-Martin et al. 2008), G1 isoform in glioblastoma (Di Cristofori, Ferrero et al. 2015), *c* subunit in hepatocellular carcinoma (Xu, Xie et al. 2012), and *E1* isoform in EC (Son, Kim et al. 2016). This study aims at shedding a new light on this controversial issue by looking at the gene expression pattern for all V-ATPase subunits in ESCC and analyzing the expression balances among the putatively isoforms distinctive in relation to normal tissues. The data obtained not only open a new perspective on the structural and functional bases of the differential activation for this proton pump in ESCC, but also provide compelling evidences for wider significance to the carcinogenesis process in general. For this purpose, we have also performed the Oncomine database analysis, which revealed that specific expression patterns of V-ATPase subunits are not restricted to esophageal cancers since a similar imbalance of isoforms found in ESCC was also detected in different cancer types (**Figure S4**). The differential expression pattern we have found in tumors, encompassing all V-ATPase subunits isoforms, might also imply in specific coupling regulation and activation patterns quite different from that exhibited by normal cells, but highly functional for the cancer specific ion signaling and energy transduction demands.

Our findings also suggest that the ratio of subunits isoforms can be part of a cellular code that controls the regulation of V-ATPase and tumorigenic related events. The data highlighted isoforms of subunits *a, C, d, G*, and the subunit *c”* as the most differentially expressed in ESCC (**Figure S2**), which in turn are subunits involved in the structural and functional coupling between the domains V1 and V0. This reinforces the idea that the active V-ATPase holoenzyme conformations in tumors and normal tissues might assume quite distinct regulatory couplings. Perhaps not surprisingly, subunits *A* (*ATP6V1A*) and *B* (*ATP6V1B1* and *ATP6V1B2*), which integrate the ATP hydrolysis catalytic domain, displayed no significant differential expression in ESCC comparing to normal esophageal tissues (**Figure S1**), suggesting a conserved and stable expression of the main subunits required for the substrate hydrolysis. Accordingly, the V-ATPase catalytic domain has been found to be highly conserved not only during molecular evolution and speciation, but also in relation to the counterpart domain of the F-ATPase synthase (Cross and Muller 2004).

We observed the most significant changes in the subunits *a* and *C*, which presented the highest differential expression among their respective isoforms not only in ESCC (**Figure 1**, **Figure S3**) but also in other cancers as revealed by the Oncomine analysis (**Figure S4**). Our data, in conjunction with other information in the literature, have revealed tumor-specific imbalances in the expression of these subunits. For instance, in our analyzes the isoforms *a3* (*TCIRG1*) and *a4* (*ATP6V0A4*) were upregulated and downregulated, respectively, in tumors as compared to normal tissues. Likewise, in ovarian cancer tissues, the *a3* isoform was found to be conspicuously expressed compared to normal samples (Kulshrestha, Katara et al. 2015). Overexpression of *a3* isoform results in the targeting of V-ATPases to the plasma membrane of breast cancer cells, and confers high invasiveness to breast cancer and melanoma cells lines (Nishisho, Hata et al. 2011, Capecci and Forgac 2013). Moreover, a knockdown of *a4* did not affect invasion of breast cancer cells, but unexpectedly resulted in a 2-fold increase in the *a3* mRNA levels (Capecci and Forgac 2013), which is in line with our results suggesting a tightly regulated balance among the isoforms of these V-ATPase subunits.

Our approach can also help to clarify the functional and evolutionary rational for the existence of such isoforms, an issue that remains poorly understood, since previous works have generally addressed only the most expressed isoforms for each subunit. This is the case of the *C1* isoform (*ATP6V1C1*), for which the overexpression (Otero-Rey, Somoza-Martin et al. 2008) and related potential as biomarker (Perez-Sayans, Reboiras-Lopez et al. 2010) have been described in oral cancer. The *C1* isoform overexpression has also been correlated with rearrangements of actin filaments in breast cancer cells (Cai, Liu et al. 2014), and its silencing prevented breast cancer growth and bone metastasis in murine model (Feng, Zhu et al. 2013). More recently, it was reported that *ATP6V1C1* enhanced breast cancer growth through activation of the mTORC1 pathway (McConnell, Feng et al. 2017). On the other hand, the *C2a* and *C2b* isoforms (*ATP6V1C2*) have been much less studied, most likely because these isoforms were initially proposed to be expressed specifically in kidney, lung and epididymis (Smith, Borthwick et al. 2002, Sun-Wada, Murata et al. 2003, Sun-Wada, Yoshimizu et al. 2003). However, we have found similar levels of expression for both *C1* and *C2* isoforms in normal esophageal surrounding tissues, contrasting with their expression imbalance in tumors characterized by a shift towards a strong *C1* dominance (**Figure 2** and **Figure S5**).

The validation of the transcriptome data by qRT-PCR reinforced that the ESCC group exhibits an increase of about 2.5-fold in *ATP6V1C1* mRNA expression and a decrease of almost 10-fold in *ATP6V1C2* expression compared with the normal surrounding tissues group. Furthermore, we demonstrated the potential of these isoforms to be used as diagnostic markers, since their mRNA levels were able to clearly discriminate ESCC from normal surrounding esophageal tissues, with the best results achieved using the *C1/C2* ratio (**Figure 2** and **3**) rather than any individual expression pattern. This highlights an interconnectivity among the *C* isoforms expression, with a wide range of *ATP6V1C1* and *ATP6V1C2* expression between the tumors of different individuals, but invariably involving a quite proportional prevalence of the *ATP6V1C1* expression over that of *ATP6V1C2*. The analysis of TCGA database confirmed that the imbalance between the *ATP6V1C1* and *ATP6V1C2* genes expression towards the prevalence of the first over the second is a clear molecular signature of esophageal tumors, regardless the histological type (**Figure 4**). It has been shown that EAC shares several molecular alterations with gastric adenocarcinoma, whilst ESCC is closer related to squamous carcinomas of head and neck (Nancarrow, Clouston et al. 2011). In fact, EAC and ESCC differ in histological type, areas of incidence around the world, risk factors and characteristic molecular alterations (Bird-Lieberman and Fitzgerald 2009, Rustgi and El-Serag 2014). Therefore, the similar expression patterns for *C* isoforms observed in both tumors provides further evidence for this distinctive imbalance of *C* isoforms as a molecular signature of the proton pump in different cancers.

This finding is also of special interest since subunit *C* contributes to the structural and regulatory assembly of V-ATPase through the reversible dissociation of V1 and V0 domains (Puopolo, Sczekan et al. 1992, Curtis, Francis et al. 2002, Smardon and Kane 2007). Interestingly, the *C* subunit is the only of the 14 subunits that is released into the cytosol during the regulatory disassembly of the proton pump (Kane and Parra 2000). In insect models, which have no alternative isoform for *C* subunit in their genomes, this single subunit binds to polymeric F-actin and monomeric G-actin and increases the initial rate of actin polymerization in a concentration-dependent manner, regulating the thickness of actin filaments cross-linking (Vitavska, Merzendorfer et al. 2005). The interaction of actin and *C* subunit might also contribute to stabilize the proton pump in its assembled state (Serra-Peinado, Sicart et al. 2016). Likewise, the cytoskeleton rearrangement somehow interconnected with the V-ATPase activity has been implicated in key cancer-related processes, such as increased migration and invasion (Cai, Liu et al. 2014), release of exosomes containing DNA, mRNA, miRNA and protein of tumors (Liegeois, Benedetto et al. 2006), membrane fusion (Strasser, Iwaszkiewicz et al. 2011), and receptors recycling (Nishi and Forgac 2002). Therefore, it seems likely that humans and other complex organisms that express multiple C isoforms would exhibit a wider array of actin-proton pump interactions and enzymatic regulation, in comparison to other organisms encoding only one type of subunit C.

To shed light on this issue, we modelled the three-dimensional structures of each *C* isoform *in silico*. The most striking difference between C1, C2b and C2a isoforms concerns an additional stretch of 46 amino acids endowed with binding sites for different kinases and phosphatases (**Figure 5**, **Tables 3** and **4**), suggesting post-translational modifications in C2a*-*assembled holoenzyme. A structural model for *C2a* isoform, in which the additional peptide stretch assumed a beta sheet folding flanked by unstructured regions was produced by Swiss-Model (Holliday 2014). Conversely, stereochemical quality analysis performed in this study highlighted an alpha-helix bend flanked by unstructured regions as the most stable structural model for such region (**Figure 5c**). To clarify this question, we simulated a docking between the predicted models for *C2a* and PKA and PKC kinases that are known to bind to the subunit *C* (Voss, Vitavska et al. 2007). The only model that allows *C2a* binding to PKA and PKC was that with the additional peptide stretch structured in a beta sheet folding flanked by unstructured regions (**Figure 5d**, **Figure S7**). It is possible that both conformations can be expressed, each one favored by different metabolic situations, especially considering that the *C* subunit is exposed to distinct microenvironments, in the membrane bound holoenzyme and free in the cytosol during the dissociation V1-V0 regulatory cycle. This mechanism also involves the interaction of the *C* and *B* subunits with actin filaments (Serra-Peinado, Sicart et al. 2016), and based on the actin-binding domain described for the *B* subunit (Holliday, Lu et al. 2000, Chen, Bubb et al. 2004, Zuo, Vergara et al. 2008), we found similar domains in the *C* subunit isoforms (**Figure S8**). Curiously, the sequence corresponding to the putative actin binding site in the *C2a* isoform was located immediately downstream the additional peptide stretch, found only in this isoform and predicted to be highly mobile, what seems quite mechanistically sound.

Nonetheless, the mRNA of *ATP6V1C2-v2* (*C2b* isoform) was found to be barely expressed in the evaluated samples, precluding its relative quantification by qPCR. It was also possible to determine a highly significant correlation between mRNA levels of *ATP6V1C2-v1* and *ATP6V1C2-v1,2* (*r* = 0.9521, p<0.00001). Therefore, it is likely that that most of the expression observed in *ATP6V1C2-v1,2* analysis is due to the expression of the variant 1 (C2a isoform). In this context, the observed switch towards *C1* in tumors assumes a major importance, as the additional peptide stretch in the *C2a* isoform may integrate kinases-dependent and actin binding mechanisms and other related regulatory functions of normal cells, which are disturbed as the *C1* isoform becomes ubiquitously prevalent in cancer cells. This hypothesis suggested by our *in silico* models should prompt future studies on the functional relevance of this *C2a* specific peptide stretch for the H^+^-pump regulatory coupling and cytoskeleton interactions.

In conclusion, to the best of our knowledge, this is the first report on multi-cancer molecular signatures for the V-ATPase holoenzyme, indicating a key role for the differential expression of their subunits and isoforms to form a conformational code, which ultimately would lead to specific pH signals and energy transductions related to carcinogenesis. Compelling evidence was also provided for a clear distinctive imbalance of the *C* subunit isoforms pointing to *ATP6V1C1* and *ATP6V1C2* as promising targets for development of new cancer biomarkers and further in-depth mechanistic characterization. Taken together these findings open a new avenue towards understanding the functions of the differential expression of V-ATPase subunits isoforms, which hopefully would lead to future breakthroughs in exploration of this proton pump mechanistic role in carcinogenesis, cancer early diagnosis and therapeutic potential.

## MATERIALS AND METHODS

### V-ATPase gene expression in ESCC Transcriptome

V-ATPase gene expression profile of ESCC data were analyzed using transcriptome data developed by the Brazilian National Institute of Cancer – INCA (Nicolau-Neto, Da Costa et al. 2018). Microarrays were performed using a genechip Human Exon 1.0 ST array (Affymetrix, Inc.) and compared 15 ESCC paired samples (ESCC and respective surrounding tissues) and 5 healthy esophageal tissues of individuals without cancer. The raw data were normalized in Expression Console software (Affymetrix) using robust multi-array average (RMA) method. The differential expression was analyzed using the “Limma” package of R software (http://www.bioconductor.org/packages/release/bioc/html/limma.html) (Diboun, Wernisch et al. 2006, Ritchie, Phipson et al. 2015).

### V-ATPase gene expression by Oncomine database

The expression levels of V-ATPase genes in 18 cancer types were obtained from the Oncomine database (https://www.oncomine.org/resource/login.html) (Rhodes, Yu et al. 2004). The fold-change of mRNA expression in cancer tissue compared to in their normal tissues was acquired using parameters of a threshold *p*-value of 1E-4; fold-change of 2; and gene ranking in the top 10%.

### Patients and samples

Thirty-eight patients enrolled from 2008 to 2015 at the Brazilian National Cancer Institute (INCA) who had confirmed ESCC diagnosis and had not undergone chemotherapy and/or radiotherapy were recruited. Paired biopsies (tumors and normal-appearing surrounding mucosa, collected at least 5 cm far from the tumor border) were included in real-time PCR analysis. Epidemiological and clinicopathological data were obtained through interviews by using a standardized questionnaire and from patient’s medical records, including whenever possible sex, age at diagnosis, tobacco smoking, alcohol intake, tumor stage and differentiation, esophageal localization and overall survival time after diagnosis. Individuals were classified regarding tobacco smoking as smokers (for ever- or ex-smoking) or never-smoking and regarding alcohol intake as drinkers (for ever- or ex-drinkers) or never-drinkers alcohol intake. The use of the human samples was approved by the Ethic Committee of the institution (INCA - 116/11). All patients, who kindly agreed to participate in the study, signed an informed consent form prior to study enrollment

### Clinicopathological features

The clinicopathological characteristics of the 38 ESCC patients evaluated in the study are presented in **Table 5**. The median age of patients was 57 years, ranging from 48 to 79 years, male patients represented 84.2% of cases and near 90% of all patients were alcohol consumers and/or smokers. Tumors were most often located in the middle third of the esophagus (47.4%), with a high prevalence of advanced stage (III or IV) of the disease (48%) and poorly or moderately differentiated (97.4%) culminating in a high mortality rate (81.6%) with median overall survival of 7.73 months.

### Real-time PCR

Total RNA was extracted from the fresh tumor and normal surrounding esophageal tissues using RNeasy mini kit (Qiagen), according to the manufacturer’s protocol. RNA concentration and purity were determined by spectrophotometry, and 500 ng of total RNA was used for reverse transcription in a final volume of 20 μL with SuperScriptTM II Reverse Transcriptase (Invitrogen^®^), following the manufacturer’s protocol. Expression levels were detected by StepOnePlus™ Real-Time PCR System (Applied) and each reaction (12 μL) contained the corresponding cDNA (1:10), 900 ng of each primer and 6.0 μL of Power SYBR Master Mix PCR (Applied). The reaction conditions were as follows: 95°C for 10 min; 95°C for 15 sec, 60°C for 1 min, for 40 cycles; at the end a melting curve analysis was included, and fluorescence was measured from 60 to 99 °C (each sample was analyzed in duplicate). Relative mRNA levels were quantified using the comparative cycle threshold method (ΔΔCt) and normalized by the *GAPDH* expression and using the normal surrounding tissue as the reference (2^−ΔΔCt^ formula). The expression ratio between *ATP6V1C1* and *ATP6V1C2* isoform C transcripts was calculated for each patient.

The oligonucleotide sequences are shown in the **Table 6**. Transcripts of the *ATP6V1C2* variant 2 could not be evaluated individually due to detection limit of real-time PCR equipment that was not able to determine quantitatively with accuracy its very low RNA concentrations (C_T_ > 32). Therefore, the transcripts of *ATP6V1C1, ATP6V1C2* variant 1 (v1) and *ATP6V1C2* variants 1 and 2 (v1,2) were evaluated together. The cDNA samples availability determined the respective number of assays, 38 paired samples for the *ATP6V1C1*, 37 for *ATP6V1C2-v1* and 36 for *ATP6V1C2-v1,2*.

### Analysis of *ATP6V1C1* and *ATP6V1C2* expression from data deposited in The Cancer Genome Atlas (TCGA) database

Gene expression data from EC samples (n=182), both esophageal adenocarcinoma (n=87) and squamous cell carcinoma (n=95), were downloaded from the public database cBioPortal for Cancer Genomics (Cerami, Gao et al. 2012, Gao, Aksoy et al. 2013), which provides visualization, analysis, and download of largescale data sets deposited in The Cancer Genome Atlas (TCGA).

### *In silico* 3D models of *C* subunit isoforms, specific peptide domains, and protein-protein interaction sites and docking assays

Conserved polypeptide sequences were estimated by using the NCBi database and the program BLASTP 2.8.0 (Boratyn, Schaffer et al. 2012), while the analysis of conserved domains was verified by using the CDD/SPARCLE (Marchler-Bauer, Bo et al. 2017). The identification of sites for phosphorylation/dephosphorylation and protein-protein interactions was performed using the *Human Protein Reference Database* (Amanchy, Periaswamy et al. 2007). The prediction of 3D models for hC1 (NP_001686.1), hC2b (NP_653184.2) and hC2a (NP_001034451.1) isoforms was performed by homology and *ab initio* modeling by four distinct programs: Modeller 9.19 (Sali and Blundell 1993), Swiss-Model (Biasini, Bienert et al. 2014), RaptorX (Kallberg, Wang et al. 2012) and I-Tasser (Yang, Yan et al. 2015). Analysis of global and local stereochemical quality for all predicted models were also performed by using different programs: Rampage (Lovell, Davis et al. 2003), Verify3D (Eisenberg, Luthy et al. 1997), ProSA (Wiederstein and Sippl 2007), VoroMQA (Olechnovic and Venclovas 2017), ProQ3D (Uziela, Menendez Hurtado et al. 2017), Qprob (Cao and Cheng 2016), DeepQA (Cao, Bhattacharya et al. 2016) and SVMQA (Manavalan and Lee 2017). The best models were refined using the ModRefiner program (Xu and Zhang 2011). Protein-protein docking assays were carried out using the programs ClusProV2 (Comeau, Gatchell et al. 2004) and HADDOCK (van Zundert, Rodrigues et al. 2016). In order to select the best interactions and to discard false positives, extensive analysis were carried out with the programs: DockScore (Malhotra, Mathew et al. 2015), PPCheck (Sukhwal and Sowdhamini 2015) and CCharPPI (Moal, Jimenez-Garcia et al. 2015). After selection of the protein complexes with native interaction characteristics, the prediction of binding free energy (Δ*G* - kcal/mol) and the dissociation constant (*K*_d_) of the complexes were calculated using the program PRODIGY (Vangone and Bonvin 2015, Xue, Rodrigues et al. 2016). Native complexes and their interactions were modeled by the program UCSF Chimera 1.11.2 (Pettersen, Goddard et al. 2004). Multiple sequence alignments were generated using ClustalW (http://www.genome.jp/tools-bin/clustalw).

### Statistical Analysis

The data were statistically analyzed using GraphPad Prism 5 Software (San Diego, CA, USA) and the difference considered significant at a *p* value of < 0.05. Unpaired *t* test or Mann-Whitney test was applied when comparing two groups. When comparing three or more groups, one-way ANOVA or Kruskal-Wallis test and Dunn’s posttest were used. A receiver operating characteristic (ROC) curve was plotted to investigate whether *C* isoforms expression could be used as a marker to discriminate ESCC from normal-appearing surrounding mucosa.

## Supporting information

## ACKNOWLEDGEMENTS

We thank all patients involved in this study, the staffs of Endoscopy Service and National Tumor Bank of the Brazilian National Cancer Institute for their contribution in sample collection.

## ADDITIONAL INFORMATION

## Author contributions

JCVCS performed most of the experiments and analysis. PNN participated in the analysis and acquisition of data. EPC performed the *in silico* structural models. ARF and LFRP coordinated the project. JCVCS and ARF wrote the manuscript. JCVCS, PNN, TAS, ARF and LFRP performed study design. TAS and PNN participated in the collection of samples. ALOF and FFF provided specialized scientific and technical support. All authors discussed the results and manuscript text. All authors read and approved the final manuscript.

## Ethics

Human subjects: The use of the human samples was approved by the Ethic Committee of the institution (INCA - 116/11). All patients, who kindly agreed to participate in the study, signed an informed consent form prior to study enrollment.

## Abbreviations

DAB: Diaminobenzidin
EAC: Esophageal adenocarcinoma
EC: Esophageal cancer
ESCC: Esophageal squamous cell carcinoma
GAPDH: Glyceraldehyde-3-phosphate dehydrogenase
TCGA: The Cancer Genome Atlas

## SUPPLEMENTARY MATERIALS

**Figure S1.** Expression levels of *A* and *B* subunits of V-ATPase in tumor, normal-appearing surrounding tissue, and healthy esophageal mucosa.

**Figure S2.** Expression levels of *c”* subunit and *d* and *G* isoforms in tumor, normal-appearing surrounding tissue and healthy esophageal mucosa.

**Figure S3.** Expression levels of *a3* and *a4* isoforms in tumor, normal-appearing surrounding tissue and healthy esophageal mucosa.

**Figure S4.** mRNA expression pattern of V-ATPase genes in several cancer types.

**FigureS5.** Distinctive balance of C isoforms.

**FigureS6.** *C* isoforms mRNA expression in esophageal cancer histological subtypes.

**Figure S7.** Protein-protein dockings between proposed hC2a models with already resolved structure of PKC and PKA proteins.

**Figure S8.** Putative actin-binding sites in C isoforms.

**Table 1.** Fold-change mean values comparing ESCC and normal surrounding tissues for each V-ATPase gene.

**Table 2.** Similarities between V-ATPase C subunits.

**Table 3.** Serine kinase/phosphatase motifs present in the additional exon of hC2a isoform.

**Table 4**. Phosphorylated serine-based motifs in the additional exon of hC2a isoform.

**Table 5.** Characteristics of the individuals included in the study.

**Table 6.** Oligonucleotide sequences and amplicon size.

