## Supplementary material for "Cancer V-ATPase Expression Signatures: A Distinctive Balance of Subunit *C* Isoforms in Esophageal Carcinoma"

**SUPPLEMENTARY FIGURES**

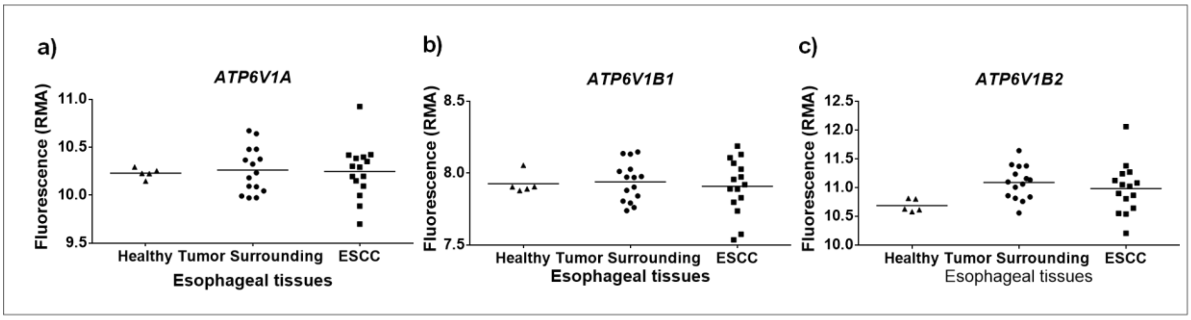


**Figure S1.** Expression levels of *A* and *B* subunits of V-ATPase in tumor, normal-appearing surrounding tissue, and healthy esophageal mucosa.

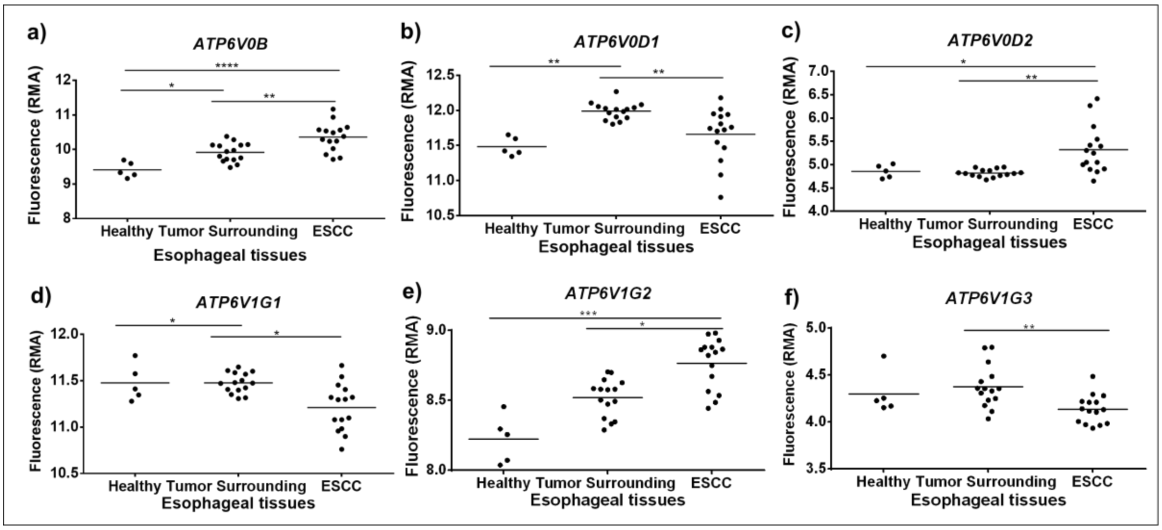


**Figure S2.** Expression levels of c" subunit and *d* and *G* isoforms in tumor, normal-appearing surrounding tissue and healthy esophageal mucosa. $*p < 0.05$; $**p < 0.01$; $***p < 0.001$; $****p < 0.0001$.

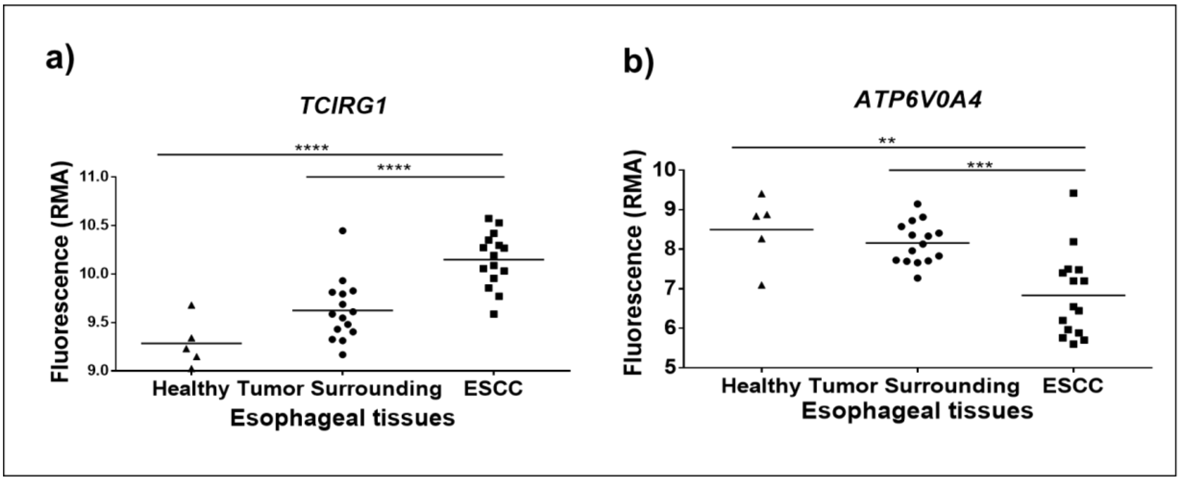


**Figure S3.** Expression levels of a3 and a4 isoforms in tumor, normal-appearing surrounding tissue and healthy esophageal mucosa. $**p<0.01$; $***p<0.001$; $****p<0.0001$.

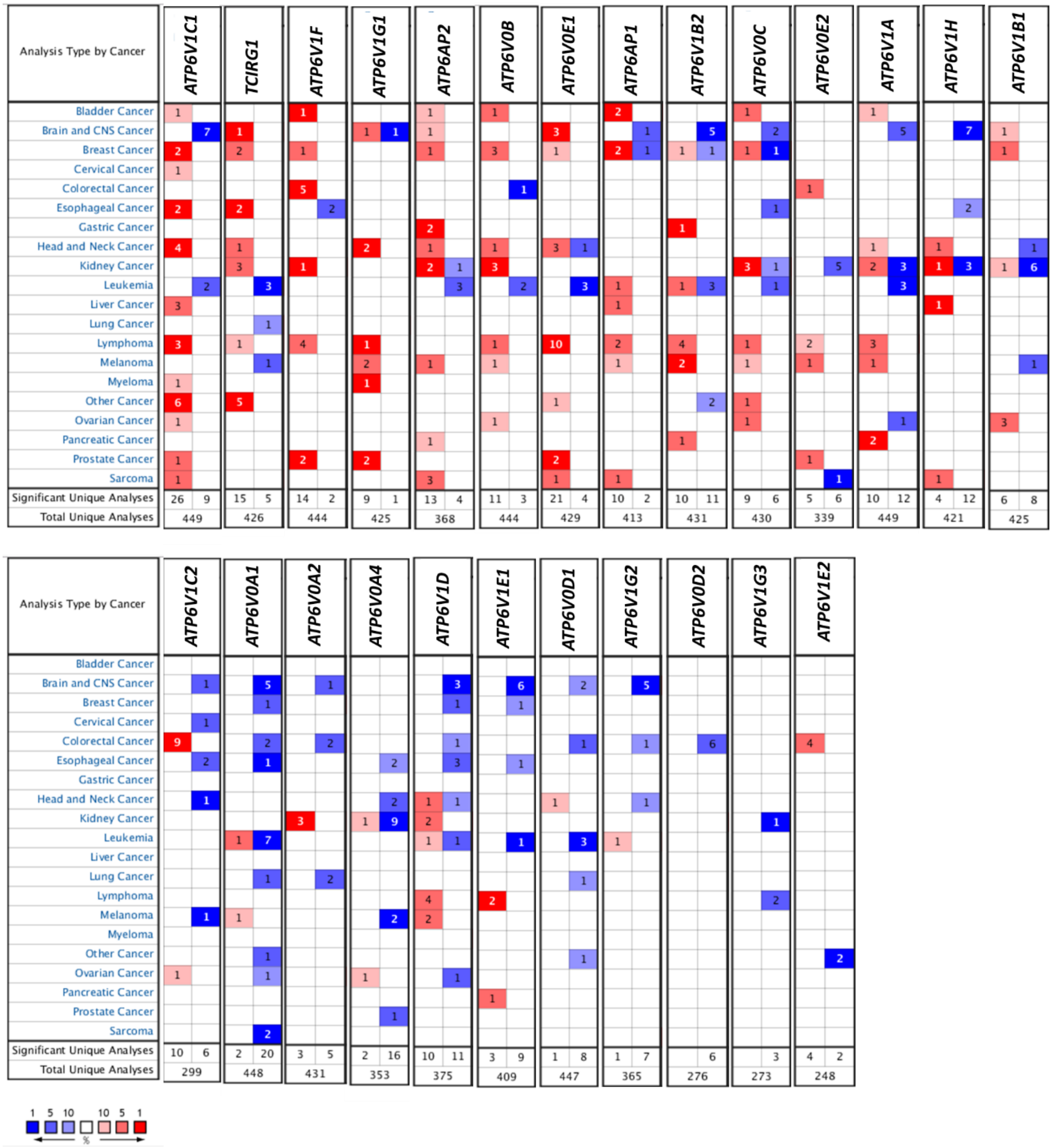


**Figure S4. mRNA expression pattern of V-ATPase genes in several cancer types**. The comparison indicates the number of the analyses that meet the threshold with mRNA over-expression (left column, red) and under-expression (right column, blue) in cancer versus corresponding normal tissues. Each column refers to a V-ATPase gene and each line refers to a cancer site (some tumor sites include different histological types). The threshold was designed with the following parameters: *p*-value of 1E-4, fold-change of 2, and gene ranking of 10%.

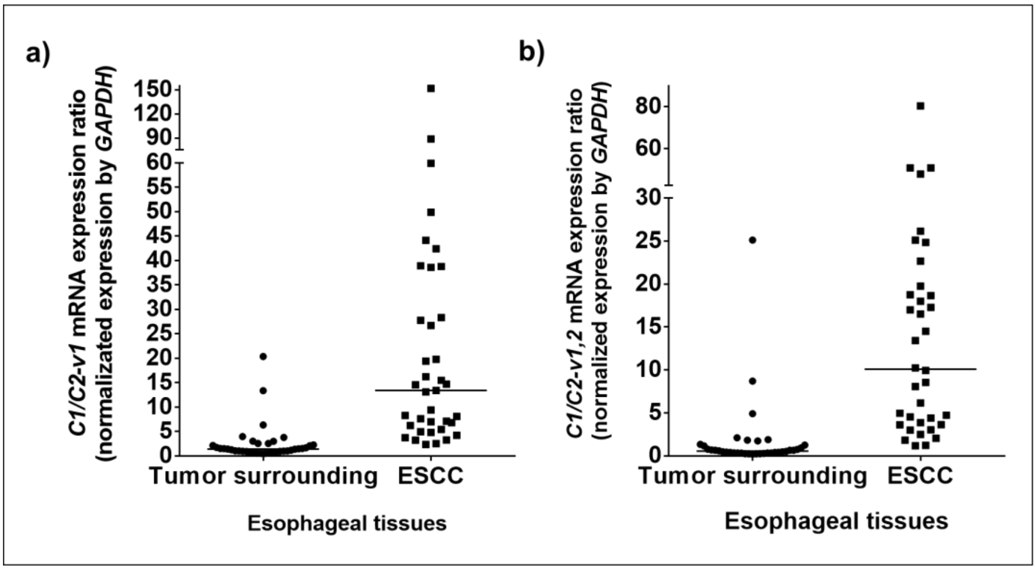


**Figure S5. Distinctive balance of C isoforms.** The ratio of expression: (a) *ATP6V1C1* by *ATP6V1C2-v1* and (b) *ATP6V1C1* by *ATP6V1C2-v1,2.* mRNA levels were normalized by those of *GAPDH*, used as the housekeeping gene. $p < 0.0001$.

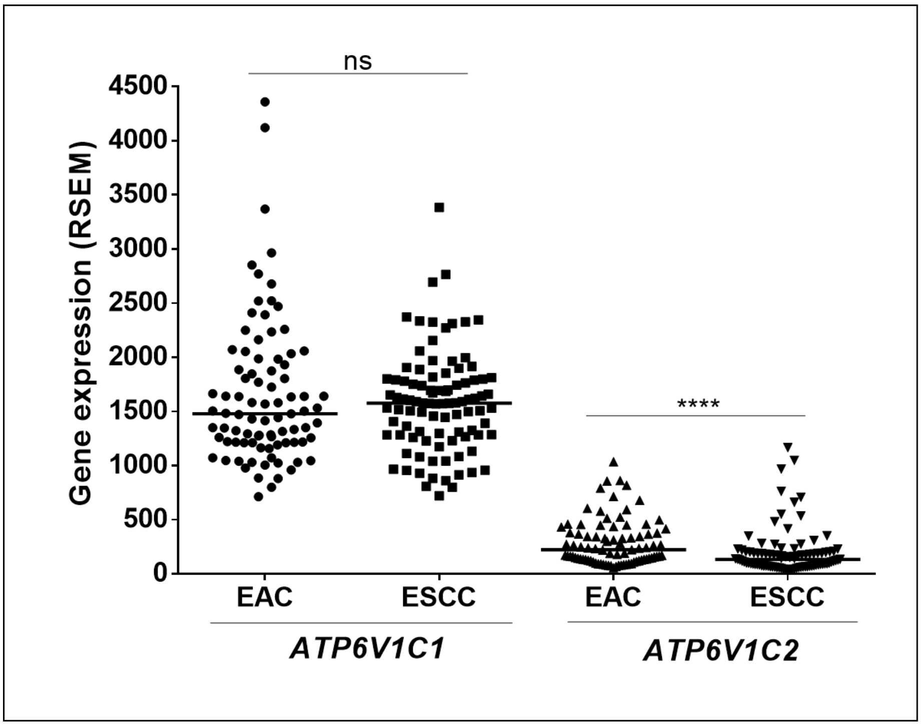


**Figure S6. *C* isoforms mRNA expression in esophageal cancer histological subtypes**. *ATP6V1C1* and *ATP6V1C2* expression median obtained in the TCGA comparing EAC and ESCC tissues. ****$p < 0.0001$. ns – not significant.

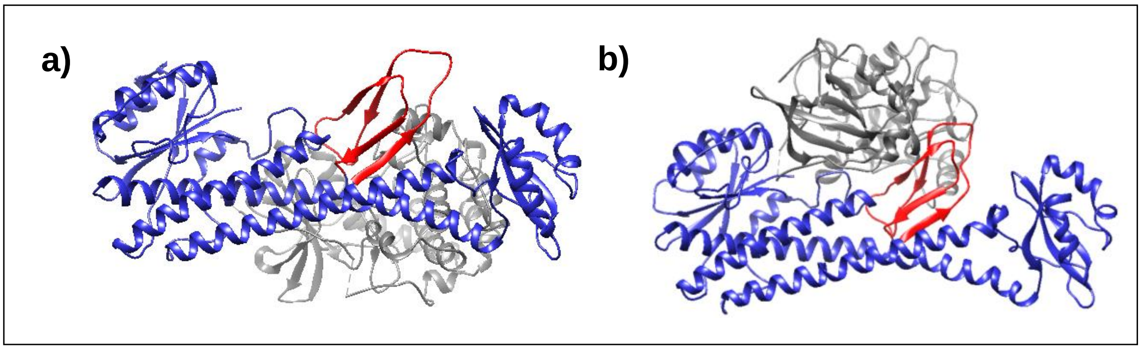


**Figure S7.** Protein-protein dockings between proposed hC2a models with already resolved structures of (a) PKA and (b) PKC.

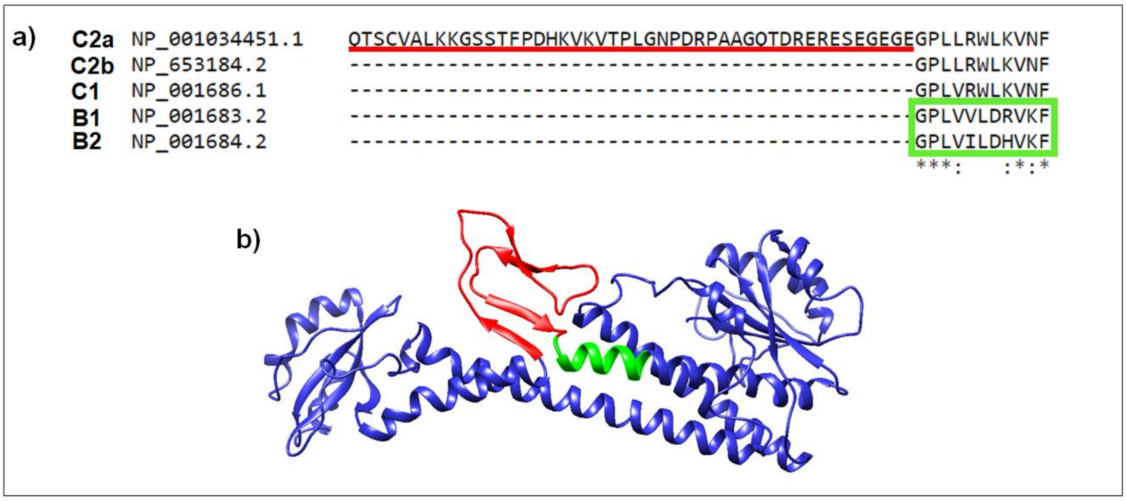


**Figure S8. Putative actin-binding sites in *C* isoforms.** (a) Multiple sequence alignment comparing “profilin-like” domain in subunit *B* (green box) and similar sequence in *C* subunit isoforms**.** (b) 3D structure of *C2a* isoform highlighting in green the putative "profilin-like" domain for actin-binding. The additional peptide stretch of 46 aa present in the hC2a isoform is shown in red.
