## Supplementary material for "Cancer V-ATPase Expression Signatures: A Distinctive Balance of Subunit *C* Isoforms in Esophageal Carcinoma"

### SUPPLEMENTARY TABLES

**Table S1.** Fold-change mean values comparing ESCC and normal surrounding tissues for each V-ATPase gene.

| Gene | Chromosome | Fold Change | P value | Cluster ID |
|---|---|---|---|---|
| *ATP6V1A1* | 3q13.31 | -1 | 0.78 | 2636589 |
| *ATP6V1B1* | 2p13.1 | -1 | 0.79 | 2487918 |
| *ATP6V1B2* | 8p21.3 | -1,08 | 0.45 | 3088544 |
| *ATP6V1C1* | 8q22.3 | 2,79 | 0.000001 | 3110171 |
| *ATP6V1C2* | 2p25.1 | -1,28 | 0.000004 | 2469575 |
| *ATP6V1D* | 14q23-q24.2 | 1,27 | 0.01 | 3569200 |
| *ATP6V1E1* | 22q11.1 | -1,28 | 0.06 | 3951887 |
| *ATP6V1E2* | 2p21 | 1,07 | 0.009 | 2551651 |
| *ATP6V1F* | 7q32 | -1,07 | 0.078 | 3023211 |
| *ATP6V1G1* | 9q32 | -1,18 | 0.0010 | 3186191 |
| *ATP6V1G2* | 6p21.3 | 1,21 | 0.0003 | 2949038 |
| *ATP6V1G3* | 1q31.3 | -1,19 | 0.001232 | 2449922 |
| *ATP6V1H* | 8q11.2 | -1,22 | 0.0479 | 3135452 |
| *ATP6V0A1* | 17q21 | -1,09 | 0.55 | 3721718 |
| *ATP6V0A2* | 12q24.31 | -1,14 | 0.003 | 3436021 |
| *TCIRG1* | 11q13.2 | 1,53 | 0.000037 | 3337390 |
| *ATP6V0A4* | 7q34 | -2,76 | 0.00015 | 3075381 |
| *ATP6V0C* | 16p13.3 | -1,1 | 0.01 | 3644887 |
| *ATP6V0B* | 1p32.3 | 1.43 | 0.0031 | 2333658 |
| *ATP6V0D1* | 16q22.1 | -1,17 | 0.00269 | 3695699 |
| *ATP6V0D2* | 8q21.3 | 1,30 | 0.0009 | 3105749 |
| *ATP6V0E1* | 5q35.1 | -1,04 | 0.02 | 2841284 |
| *ATP6V0E2* | 7q36.1 | -1,07 | 0.1485 | 3031181 |
| *ATP6AP1* | Xq28 | -1,20 | 0.019 | 3996381 |
| *ATP6AP2* | Xp11.4 | 1,12 | 0.254 | 3974556 |

* Values were obtained from a previously performed microarray analysis.

**Table S2.** Similarities between V-ATPase C subunits.

| | ID/Sim (%) | | | R.M.S.D. | | |
|---|---|---|---|---|---|---|
| | **C1** | **C2a** | **C2b** | **C1** | **C2a** | **C2b** |
| **C1** | --- | 55.3/74.2 | 61.9/83.1 | --- | 1.920 | 1.493 |
| **C2a** | 55.3/74.2 | --- | 89.2/89.2 | 1.920 | --- | 2.462 |
| **C2b** | 61.9/83.1 | 89.2/89.2 | --- | 1.493 | 2.462 | --- |

ID – identity; Sim – similarity; R.M.S.D. - Root Mean Square deviation.

**Table S3.** Serine kinase/phosphatase motifs present in the additional exon of hC2a isoform.

| | Amino acid position | Amino acid Sequence | Corresponding motif described in the literature | Features of motif |
|---|---|---|---|---|
| 1 | 1 - 6 | QTSCVA | X[pS/pT]XXX[A/P/S/T] | G protein-coupled receptor kinase 1 substrate |
| 2 | 7 - 12 | LKKGSS | [M/I/L/V]X[R/K]XX[pS/pT] | Chk1 kinase substrate motif |
| 3 | 7 - 14 | LKKGSSTF | [M/V/L/I/F]X[R/K]XX[pS/pT]XX | Calmodulin-dependent protein kinase II substrate motif |
| 4 | 8 - 11 | KKGS | KXX[pS/pT] | PKA kinase substrate motif |
| 5 | 8 - 11 | KKGS | [R/K]XX[pS/pT] | PKC kinase substrate motif |
| 6 | 8 - 11 | KKGS | [R/K][R/K]X[pS/pT] | PKA kinase substrate motif |
| 7 | 8 - 12 | KKGSS | KXXX[pS/pT] | PKA kinase substrate motif |
| 8 | 9 - 11 | KGS | [R/K]X[pS/pT] | PKA kinase substrate motif |
| 9 | 9 - 11 | KGS | [R/K]X[pS/pT] | PKC kinase substrate motif |
| 10 | 10 - 15 | GSSTFP | X[pS/pT]XXX[A/P/S/T] | G protein-coupled receptor kinase 1 substrate motif |
| 11 | 11 - 13 | SST | pSX[E/pS*/pT*] | Casein Kinase II substrate motif |
| 12 | 13 - 16 | TFPD | [pS/pT]XX[E/D] | Casein Kinase II substrate motif |
| 13 | 13 - 16 | TFPD | [pS/pT]XX[E/D/pS*/pY*] | Casein Kinase II substrate motif |
| 14 | 13 - 16 | TFPD | [pS/pT]XX[E/D] | Casein Kinase II substrate motif |
| 15 | 18 - 22 | KVKVT | KXXX[pS/pT] | PKA kinase substrate motif |
| 16 | 20 - 22 | KVT | [R/K]X[pS/pT] | PKA kinase substrate motif |
| 17 | 20 - 22 | KVT | [R/K]X[pS/pT] | PKC kinase substrate motif |
| 18 | 21 - 23 | VTP | X[pS/pT]P | GSK-3, ERK1, ERK2, CDK5 substrate motif |
| 19 | 35 - 37 | TDR | [pS/pT]X[R/K] | PKA kinase substrate motif |
| 20 | 35 - 37 | TDR | [pS/pT]X[R/K] | PKC kinase substrate motif |
| 21 | 35 - 38 | TDRE | [pS/pT]XX[E/D] | Casein Kinase II substrate motif |
| 22 | 35 - 38 | TDRE | [pS/pT]XX[E/D/pS*/pY*] | Casein Kinase II substrate motif |
| 23 | 35 - 38 | TDRE | [pS/pT]XX[E/D] | Casein Kinase II substrate motif |
| 24 | 38 - 41 | ERES | [E/D]XX[pS/pT] | Casein Kinase I substrate motif |
| 25 | 39 - 41 | RES | RXpS | PKA kinase substrate motif |
| 26 | 39 - 41 | RES | [R/K]X[pS/pT] | PKA kinase substrate motif |
| 27 | 39 - 41 | RES | [R/K]X[pS/pT] | PKC kinase substrate motif |
| 28 | 39 - 42 | RESE | XX[pS/pT]E | G protein-coupled receptor kinase 1 substrate motif |
| 29 | 40 - 44 | ESEGE | [E/D]pS[E/D]X[E/D] | Casein Kinase II substrate motif |
| 30 | 40 - 44 | ESEGE | [E/D][pS/pT]XXX | b-Adrenergic Receptor kinase substrate motif |
| 31 | 41 - 44 | SEGE | pSXX[E/D] | Casein kinase II substrate motif |
| 32 | 41 - 44 | SEGE | pSXX[E/pS*/pT*] | Casein Kinase II substrate motif |
| 33 | 41 - 44 | SEGE | [pS/pT]XX[E/D] | Casein Kinase II substrate motif |
| 34 | 41 - 44 | SEGE | [pS/pT]XX[E/D/pS*/pY*] | Casein Kinase II substrate motif |
| 35 | 41 - 44 | SEGE | [pS/pT]XX[E/D] | Casein Kinase II substrate motif |
| 36 | 41 - 46 | SEGEGE | pS[E/D]X[E/D]X[E/D] | Casein Kinase II substrate motif |

pS, pT - phosphorylated residues; * indicates the residue that must be phosphorylated for the enzyme to recognize the motif.

**Table S4**. Phosphorylated serine-based motifs in the additional exon of hC2a isoform.

| | Amino acid position | Amino acid sequence | Corresponding motif described in the literature | Features of motif |
|---|---|---|---|---|
| 1 | 11 - 13 | SST | S[pS/pT]X | MDC1 BRCT domain binding motif |
| 2 | 11 - 13 | SST | S[pS/pT]X | Plk1 PBD domain binding motif |
| 3 | 22 - 23 | TP | [pS/pT]P | WW domain binding motif |

pS, pT - phosphorylated residues.

**Table S5.** Characteristics of the individuals included in the study.

| Clinicopathological features | N | (%) |
|---|---|---|
| **Gender** | | |
| Male | 32 | 84.2% |
| Female | 6 | 10.5% |
| **Age (years)** | **Median** | **Range** |
| Male | 57 | (48-75) |
| Female | 57 | (49-79) |
| Total | 57 | (48-79) |
| **Alcohol Consumption** | **N** | **(%)** |
| No | 2 | 11.8% |
| Drinker/Ex drinker | 15 | 88.2% |
| N/A | 21 | |
| **Tobacco Consumption** | | |
| No | 1 | 5.9% |
| Smoker/Ex smoker | 16 | 94.1% |
| N/A | 21 | |
| **Tumor Site** | | |
| Upper Third | 6 | 15.8% |
| Medium Third | 18 | 47.4% |
| Lower Third | 5 | 13.2% |
| Upper-middle thirds | 5 | 13.2% |
| Middle-lower thirds | 3 | 7.9% |
| Upper-middle-lower thirds | 1 | 2.6% |
| **Tumor Stage** | | |
| I | 0 | 0.0% |
| II | 14 | 51.9% |
| III | 6 | 22.2% |
| IV | 7 | 25.9% |
| N/A | 11 | |
| **Tumor differentiation** | | |
| Well | 1 | 2.6% |
| Moderate | 30 | 78.9% |
| Poor | 7 | 18.4% |
| **Metastasis** | | |
| Yes | 4 | 22.2% |
| No | 14 | 77.8% |
| N/A | 20 | |
| **Death** | | |
| Yes | 31 | 81.6% |
| No | 7 | 18.4% |
| **Total** | 38 | 100% |

N/A = not informed.

**Table S6.** Oligonucleotide sequences and amplicon size.

| Genes | Oligonucleotide sequences (5'-3') | Amplicon size |
|---|---|---|
| ***GAPDH Var. 1-4*** | F: CAACAGCCTCAAGATCATCAGCAA<br>R: AGTGATGGCATGGACTGTGGTCAT | 124 |
| ***ATP6V1C1*** | F: ACCTGTCAGCAAACATGGGA<br>R: ACATCCAACGTGCCAACCTTT | 110 |
| ***ATP6V1C2 Var. 1*** | F: ACAGTATCAAACTTCCTGTGTTGC<br>R: ATCAGGGTTACCTAGCGGGG | 91 |
| ***ATP6V1C2 Var. 1-2*** | F: CCTGACTTCAAGGTGGGGAC<br>R: CATGACTTCCACCACGCTCT | 120 |

Var - Splicing Variant; F - forward; R - reverse; bp - base pairs.
